# Springs vs. motors: Ideal assistance in the lower limbs during walking at different speeds

**DOI:** 10.1101/2024.01.18.576164

**Authors:** Israel Luis, Maarten Afschrift, Elena M. Gutierrez-Farewik

## Abstract

Recent years have witnessed breakthroughs in assistive exoskeletons; both passive and active devices have reduced metabolic costs near preferred walking speed by assisting muscle actions. Metabolic reductions at multiple speeds should thus also be attainable. Musculoskeletal simulation can potentially predict the interaction between assistive moments, muscle-tendon mechanics, and walking energetics. In this study, we simulated devices’ optimal assistive moments based on minimal muscle activations during walking with prescribed kinematics and dynamics. We used a generic musculoskeletal model with calibrated muscle-tendon parameters and computed metabolic rates from muscle actions. We then simulated walking across multiple speeds and with two ideal actuation modes – motor-based and spring-based – to assist ankle plantarflexion, knee extension, hip flexion, and hip abduction and compared computed metabolic rates. We found that both actuation modes considerably reduced physiological joint moments but did not always reduce metabolic rates. Compared to unassisted conditions, motor-based ankle plantarflexion and hip flexion assistance reduced metabolic rates, and this effect was more pronounced as walking speed increased. Spring-based hip flexion and abduction assistance increased metabolic rates at some walking speeds despite a moderate decrease in some muscle activations. Both modes of knee extension assistance reduced metabolic rates to a small extent, even though the actuation contributed with practically the entire net knee extension moment during stance. Motor-based hip abduction assistance reduced metabolic rates more than spring-based assistance, though this reduction was relatively small. Future work should experimentally validate the effects of assistive moments and refine modeling assumptions accordingly. Our computational workflow is freely available online.

**Author Summary:** We used simulation to identify ideal assistance at major lower limb joints that can potentially be produced by motor-based or spring-based assistive devices in slow, normal, and fast walking. We found that assistance from both actuation modes decreased muscle activations and net muscle moments to varying extents, depending on joint and walking speed, but they did not always reduce metabolic energy of muscles. Motor-based assistance was overall more effective than spring-based assistance, and spring-based assistance at times increased the metabolic energy. The largest metabolic energy reductions occurred with motor-based plantarflexion assistance, followed by motor-based hip flexion assistance, both more notably at higher speeds. Motor-based hip abduction assistance also reduced metabolic energy, somewhat inversely with walking speed. Spring-based assistance was overall less effective than motor-based assistance but did reduce metabolic energy with plantarflexion assistance in slow walking and with hip flexion assistance in fast walking. Knee extension assistance, regardless of actuation mode or walking speed, had little to no influence on metabolic energy. Our simulation findings do not support knee extension assistance at all, nor spring-based hip flexion assistance in slow walking or hip abduction assistance at any speed if a device goal is to reduce muscle activations.

## 1. Introduction

Multiple lower limb exoskeletons have made breakthroughs in the past decade by improving walking and running efficiency (1). Increasingly efficient actuators and batteries, better strategies for human-device control, and lighter structures and physical interfaces have continuously improved assistance efficiency (2). Current efforts to bridge the gap between laboratory-based observations and real-world benefits focus frequently on refining methods to identify optimal assistance (3), integrating human movement intention into exoskeleton control (4), and expanding exoskeleton use to make them suitable across multiple locomotion modes (5). In this regard, musculoskeletal simulations can complement these efforts by guiding hypotheses about muscle-device interaction and revealing causal relationships in experimental observations (6).

Prior musculoskeletal simulation studies of exoskeleton assistance have provided insights into muscle-tendon mechanics and energetics, though simulation findings have not always agreed with experimental observations. Researchers have, through simulations, estimated the influence of exoskeleton assistance on tendon energy storage and release (7), on muscle fiber operating lengths and velocities (8), and on muscle activations, all of which influence muscle energetics and metabolic rates (9–11). In theory, a model-based approach can be used to design exoskeleton controllers that result in optimal muscle dynamics and minimal energy cost. For instance, Franks et al. (12) used simulations with prescribed kinematics and dynamics to predict optimal multi-joint assistive moments, i.e. leading to minimal metabolic rates during walking. In subsequent experiments with these assistive moments, they indeed observed reduced muscle excitations and metabolic costs, but not as much as the model predicted. Uchida et al. (11) used a similar computational approach to predict optimal assistive moments for running; these were later evaluated experimentally by Lee et al. (13), who reported decreased metabolic cost, but again not as much as the model predicted. Some discrepancies between simulations and experiments are to be expected, as modeling approaches rely on a number of assumptions, including simplified muscle control and dynamics, simplified or no user-device interaction forces, massless devices, and unchanged kinematics. Model-based approaches thus have potential use in informing the design of assistive interventions, more so if they can accurately estimate muscle energetics and metabolic rates.

Most musculoskeletal modeling studies aiming to predict optimal assistive moments have focused on gait at or near preferred walking speed, even though daily activities encompass a wide range of speeds and locomotion modes. Several studies have predicted optimal lower limb exoskeleton assistive moments near preferred walking speed in normal and loaded conditions, such as carrying extra weight (6)(9). Other activities, such as walking at various speeds or stair ascent, have been studied less (6). To the best of our knowledge, only two musculoskeletal-based studies have examined optimal assistance during gait at a range of speeds; Uchida et al. examined mechanics and energetics in young adults with ideal actuators during running at 2 and 5 m/s (11), and Cseke et al., in elderly adults during walking at self-selected slow (0.86 m/s), comfortable (1.22 m/s) and fast (1.53 m/s) speeds (10).

Whereas variable assistive torque profiles that can theoretically be provided by a motor can be expected to reduce metabolic rates, spring-based actuation, i.e., a spring or elastic component that can store potential energy during elongation and then release it during shortening, can also potentially influence muscle dynamics and gait energetics. Spring-based exoskeletons can also potentially be lighter and less cumbersome than motorized exoskeletons. Spring-based assistance should theoretically be effective during gait phases characterized by joint power absorption followed by joint power generation, which is the case with ankle dorsi-/plantarflexion and hip flex-/extension in pre-swing, hip ab-/adduction during midstance, and knee flex-/extension during loading response and midstance (14). Spring-based actuators have been observed experimentally to substantially decrease physiological joint moments, i.e., joint moments from muscle actions, but with negligible metabolic reduction (15). Prior musculoskeletal simulation studies have provided insights into the causal relationship between muscle mechanics and spring-based assistive moments near preferred speed (16,17) predictions have agreed with experimental observations to some degree (18,19). Simulations that investigate the influence of spring-based assistance on muscle energetics can potentially inform device design.

The objectives of the study were thus to simulate how two modes of assistance, spring-based and motor-based, at individual lower limb joints affect computed muscle dynamics and metabolic rates during walking at various speeds. We hypothesized that assistive moments will reduce muscle activations and metabolic rates and that motor-based actuation will be more efficient than spring-based actuation. Also, assisting ankle plantarflexion with any mode of assistance will yield the largest reduction of metabolic rates compared to unassisted conditions among all the muscle groups and at all walking speeds.

## 2. Methods

### 2.1 Musculoskeletal simulation workflow

We implemented a simulation workflow to estimate muscle dynamics and metabolic rates during walking using musculoskeletal models with calibrated muscle-tendon parameters and recorded motion. We used previously reported experimental data (20) and the OpenSim software (21) to scale a generic musculoskeletal model and compute joint kinematics and dynamics. We calibrated muscle-tendon parameters in the scaled musculoskeletal model to better represent fiber lengths and passive angle-moment relationships (22). Then, we performed musculoskeletal simulations while walking with prescribed kinematics and dynamics using trajectory optimization (23) (Fig. 1).

**Fig. 1.**
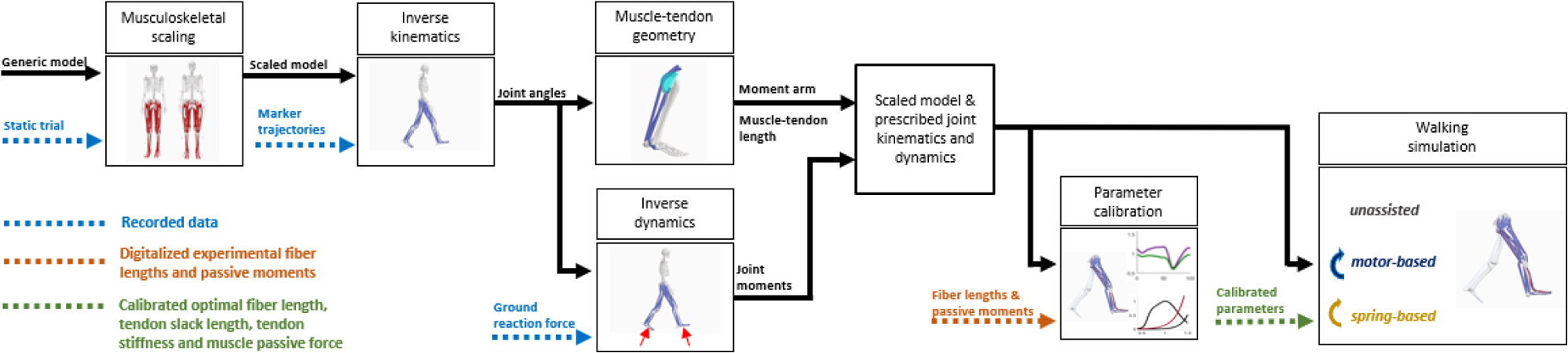
Simulation workflow per each subject. Inverse kinematics and dynamics are computed using the OpenSim workflow. Moment arms and muscle-tendon lengths are computed from the inverse kinematic solution using the Muscle Analysis tool from OpenSim. We then tuned the muscle-tendon parameters – optimal fiber length, tendon slack length, and tendon stiffness – such that the simulated muscle fiber lengths and excursions matched reported findings from ultrasound imaging. Next, we tuned the muscle passive force curves such that the simulated passive moments matched passive angle-moment joint relationships reported in an *in vivo* study. Finally, we simulated walking across various speeds with no actuators and with the various assistive actuators.

#### Experimental data

Experimental motion data: marker trajectories and ground reaction force of five unimpaired (2/3 male/female, [mean ± SD] age: 31.4 ± 7.4 years old, height: 1.75 ± 0.03 m, body mass: 69.0 ± 10.3 kg) reported in a previous publication were used for this study (20). In brief, subjects walked on a treadmill at a range of walking speeds, specifically 55%, 70%, 85%, 100%, 115%, 130%, and 145% of their estimated preferred walking speed (PWS). Subjects then walked along a lab pathway and emulated different walking speeds by matching recorded cadences from treadmill walking. Marker positions (100 Hz), based on the Conventional Gait Model with the Extended-foot model (CGM 2.4) and ground reaction forces (1000 Hz) were measured using optical motion capture (Vicon V16, Oxford, UK) and strain gauge force platforms (AMTI, Watertown, MA, USA), respectively.

#### Musculoskeletal model, joint kinematics, and inverse dynamics

A generic musculoskeletal model developed by Rajagopal et al. (24) with modified hip abductor muscle paths (25) was selected for this study. We scaled the generic model using OpenSim’s Scale Tool, which adjusted muscle paths, skeletal geometry, and segment inertial properties to fit anthropometric dimensions obtained from a captured static calibration trial. We adjusted the maximum isometric force of the soleus, gastrocnemius, and tibialis anterior as per Arnold et al. (26).

Marker trajectories and ground reaction forces throughout three gait cycles per subject at low (55% PWS), normal (100% PWS), and fast (145% PWS) walking speeds were analyzed with inverse kinematics and inverse dynamics using OpenSim 4.1. Marker tracking weights for inverse kinematics were selected to minimize the error between experimental and virtual markers. The subtalar and metatarsal joints were fixed at neutral anatomical positions.

#### Tuning of muscle-tendon parameters

We used a computational tool to tune muscle-tendon parameters such that each subject’s muscle excitations, fiber lengths, and passive moments best resembled experimental observations (22). The tuning was done in two steps. First, we tuned optimal fiber lengths, tendon slack lengths, and tendon stiffnesses of the gastrocnemius lateralis, gastrocnemius medialis, soleus, and vasti (lateralis, medialis, and intermedius) to match muscle fiber lengths and excursions obtained from those reported in from ultrasound imaging (27,28). Then, we tuned muscle passive force-length curves to match the reported passive moment at various ankle, knee, and hip joint angles from an *in vivo* study (29). Compared to simulation with the original generic model, these steps result in estimated muscle excitations that better agree with observed electromyography signals (22).

#### Solving muscle redundancy

Our implementation is based on the simulation framework proposed by De Groote et al. (23), which uses direct collocation dynamic optimization and implicitly incorporates activation and contraction dynamics. Muscle excitations, states, and state derivatives were computed based on the assumption that the muscle redundancy is solved by the optimization criterion of minimal muscle activations squared. Two other terms are present in the objective function to improve the feasibility and convergence of the formulation. Reserve actuators in each degree of freedom are added to guarantee the problem’s feasibility, and their use is penalized in the objective function. Also, a term that minimizes muscle fiber velocities is included to improve numerical computation (L2 regularization). The objective function is implemented as (1):

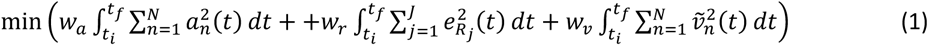

Where *a*_*n*_ is muscle activation of muscle 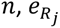 is the excitation of the reserve actuator of joint *j*, *ṽ*_*n*_ is the normalized fiber velocity of muscle *n*, *t*_*f*_ and *t*_*i*_ are the initial and final times of the gait cycle, respectively; *N* and *J* are the total number of muscles and joints in the musculoskeletal model, respectively; and *w*_*a*_, *w*_*r*_ and *w*_*v*_ are the weights of the terms in the objective function related to the muscle activations, reserve actuators, and fiber velocities, respectively. The sum of the moments produced by muscle-tendon and reserve actuators equals the net joint moment obtained from inverse dynamics at each joint. This condition was implemented as a constraint in the optimization problem as in (2)

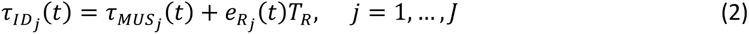

Where, at joint *j*, 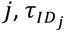 is the net joint moment, 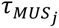 is the moment produced by the muscle-tendon actuators, subsequently referred to here as “muscle moments”, and *T*_*R*_ is the magnitude of the reserve actuator. ((Thinking about Svein’s comments, maybe comment about inertia components in the solver?))

### 2.2 Determining optimal assistive moments

Optimal assistive moments were computed using scaled musculoskeletal models with tuned muscle-tendon parameters, inverse kinematics, and inverse dynamics solutions as described above. Constraints and design variables were added when solving the muscle redundancy to model assistive moments at various muscle groups. For each mode of actuation, ideal assistive moments were added at each simulated gait cycle to assist specific muscle groups individually: plantarflexion, knee extension, hip flexion, and hip abduction. Per each subject, nine gait cycles were simulated (three gait cycles per walking speed); therefore, nine assistive moments per mode of actuation and muscle group were determined.

When solving the muscle redundancy, the objective function was the same as in unassisted conditions but constraints were added to model assistive device moment to assist muscle-tendon actuators in reproducing the inverse dynamics, i.e., the sum of the muscle moment, reserve actuator moment, and assistive device moment equals the net joint moment at each joint, as in (3)

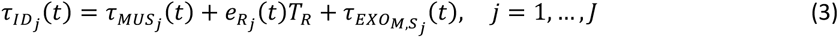

Where 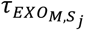 is the assistive device moment at joint *j*. In this regard, the assistive moments are the optimal solutions to assist muscles based on minimal summed muscle activations squared.

#### Motor-based moment profiles

The motor-based actuation was modeled as a unidirectional ideal moment at the corresponding degree of freedom in the musculoskeletal model. The assistive moment 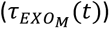 was implemented as a time-series design variable. Its magnitude was constrained to assist the aimed muscle group explicitly. For instance, 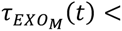 for assisting ankle plantarflexion corresponds to ankle plantarflexion moment (agonist muscle group) and avoids generating ankle dorsiflexion moment (antagonist muscle group). The motor-based actuation was not constrained in its trajectory; hence, it could have any value at each point in time to assist a muscle group. Optimal assistive moment based on motor-based actuation was determined individually for each subject and gait cycle.

#### Spring-based device parameters

The spring-based actuator was modeled as a unidirectional torsional spring that engages and disengages in specific joint angles. To implement this, we introduced three design variables: engaged timing (*t*_*c*_), disengaged timing (*t*_*d*_), and spring stiffness (*k*_*r*_), we added a constraint to impose that the angle at which the spring engaged and disengaged are similar as in (4)

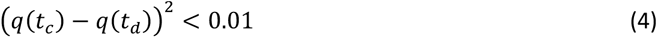

Where *q*(*t*) is the joint angle corresponding to the assisted muscle group. The assistive moment was computed as a product of the spring stiffness and the angle within the period that the spring is engaged as in (5)

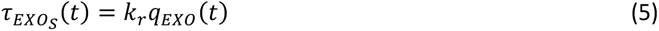

Where *q*_*EXO*_(*t*) is the joint angle displacement from the angle of engagement. This angle was modeled using hyperbolic tangent function as in (6), (7), (8), and (9)

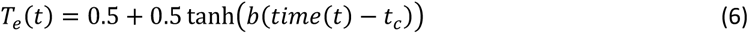

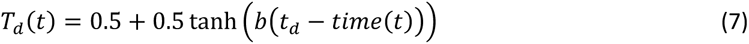

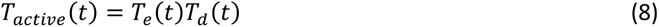

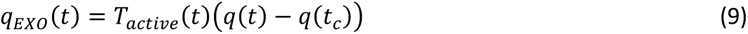

Where *T*_*e*_ is the start of the engaged period, *T*_*d*_ is the start of the disengagement period, and *T*_*active*_ (*t*) is the period where spring is engaged.

We selected *w*_*a*_, *w*_*r*_ and *w*_*v*_ as 1, 1000, and 0.001; thus, the use of reserve actuators was heavily penalized, and the influence of fiber velocities was relatively small. Also, we selected *T*_*R*_ as 100 Nm, and b as 1000 since it provided a smooth yet steep transition between null to assistive moment generation (Supplementary Fig. 1). Optimal assistive moment based on spring-based actuation was determined individually for each subject and gait cycle.

### 2.3 Metabolic rate computation

For each subject/gait cycle, each speed, and each device, the metabolic rate of each muscle was computed based on the muscle excitations, states, and state derivatives obtained from our optimization routine using a metabolic energy model proposed by Bhargava et al. (30), which we previously reported to agree with recorded metabolic rates obtained from spiroergonometry (20). In brief, muscle metabolic rate is computed as in (10)

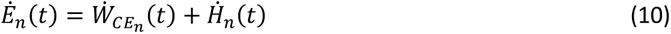

Where *Ė*_*n*_, 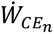 and *Ḣ*_*n*_ are the metabolic rate, contractile element work rate, and heat rate, respectively, of muscle *n*. The contractile element work rate, also called muscle power, is computed as in (11)

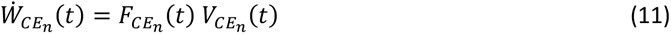

Where 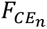 and 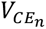 are the muscle force and fiber velocity, respectively, of muscle *n*. The heat rate depends on muscle mass, muscle activations, fiber velocities, and a function that approximates the size principle of motor recruitment, explained in detail by Bhargava et al. (30). The original formulation did not explicitly address negative metabolic rates, which are possible during eccentric contractions if muscle negative power exceeds the heat rate. As a negative metabolic rate is physiologically questionable, we adjusted it in such cases by updating the heat rate and re-computing the metabolic rate as in (12) and (13)

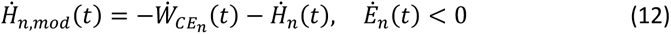

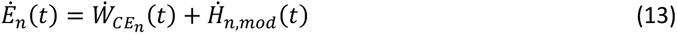

The metabolic rates for one leg (*Ė*_*L*_) is computed as the sum of all the individual muscle metabolic rates as in (14)

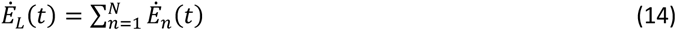

#### F. Data and statistical analysis

We evaluated the change of muscle activations, physiological joint moments, and metabolic rates between unassisted and assisted with two actuation modes during walking across speeds. Muscle activations were the sum of all the muscle activations in one leg obtained from solving the muscle redundancy and divided by the number of muscles. Net muscle moments are defined here as the net joint moments minus the assistive moments (3). Net muscle moments for agonists (plantarflexion, knee extension, hip flexion, and hip abduction) and antagonists (dorsiflexion, knee flexion, hip extension, and hip adduction) were computed in correspondence to the muscle group assisted, e.g., with ideal plantarflexion assistive moments, plantarflexion and dorsiflexion net muscle moments were presented. Metabolic rates were the sum of all the muscle metabolic rates in one leg (13). Average muscle activations, agonist and antagonist net muscle moment, and metabolic rates in unassisted and assisted conditions over each gait cycle were computed as the integral of its corresponding time-series divided by the gait cycle duration as in (15)

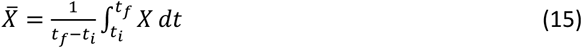

Where *X̅* are the average values of the muscle activations, agonist and antagonist net muscle moment, and metabolic rates over a gait cycle. To facilitate comparison, we computed the change (Δ) in the average muscle activations, net muscle moment, and metabolic rates for each gait cycle between unassisted and assisted conditions, and presented it as a percentage of that value in unassisted conditions as in (16)

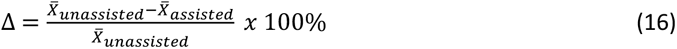

Where *X̅*_*unassisted*_ and *X̅*_*assisted*_ are the average values of the muscle activations, agonist and antagonist net muscle moment, and metabolic rates over the gait cycle in unassisted and assisted conditions, respectively. For each walking speed, we computed the average values for all subjects and gait cycles and presented the change in average metabolic rates vs. change in average muscle activations, as well as the time-series of muscle activations, agonist and antagonist net muscle moment, and metabolic rates between unassisted and assisted walking at slow (55% PWS), normal (100% PWS), and fast walking speeds (145% PWS). In addition, to complement the description of the estimated muscle-tendon mechanics and energetics, we presented the activations, work rates (obtained from (11)), and metabolic rates of individual muscles for unassisted and assisted conditions at normal walking speed in the supplementary material (average values among all subjects and gait cycles).

## 3. Results

### 3.1. Influence of assistive moments on relative muscle activations and metabolic rates

Compared to unassisted conditions, with either actuation mode, relative muscle activation changes varied, depending on the joint and muscle group assisted and with walking speed (Fig. 2). With motor-based actuation, muscle activation reduced most overall with hip flexion assistance at a high walking speed; this change decreased with decreasing walking speeds. The next highest muscle activation reduction was observed with hip abduction assistance, which, in contrast to hip flexion assistance, was proportionally higher as walking speed decreased. Muscle activations were reduced moderately with plantarflexion assistance, with a small relation to walking speed. Muscle activations were nearly unchanged with knee extension assistance at any walking speed.

**Fig. 2.**
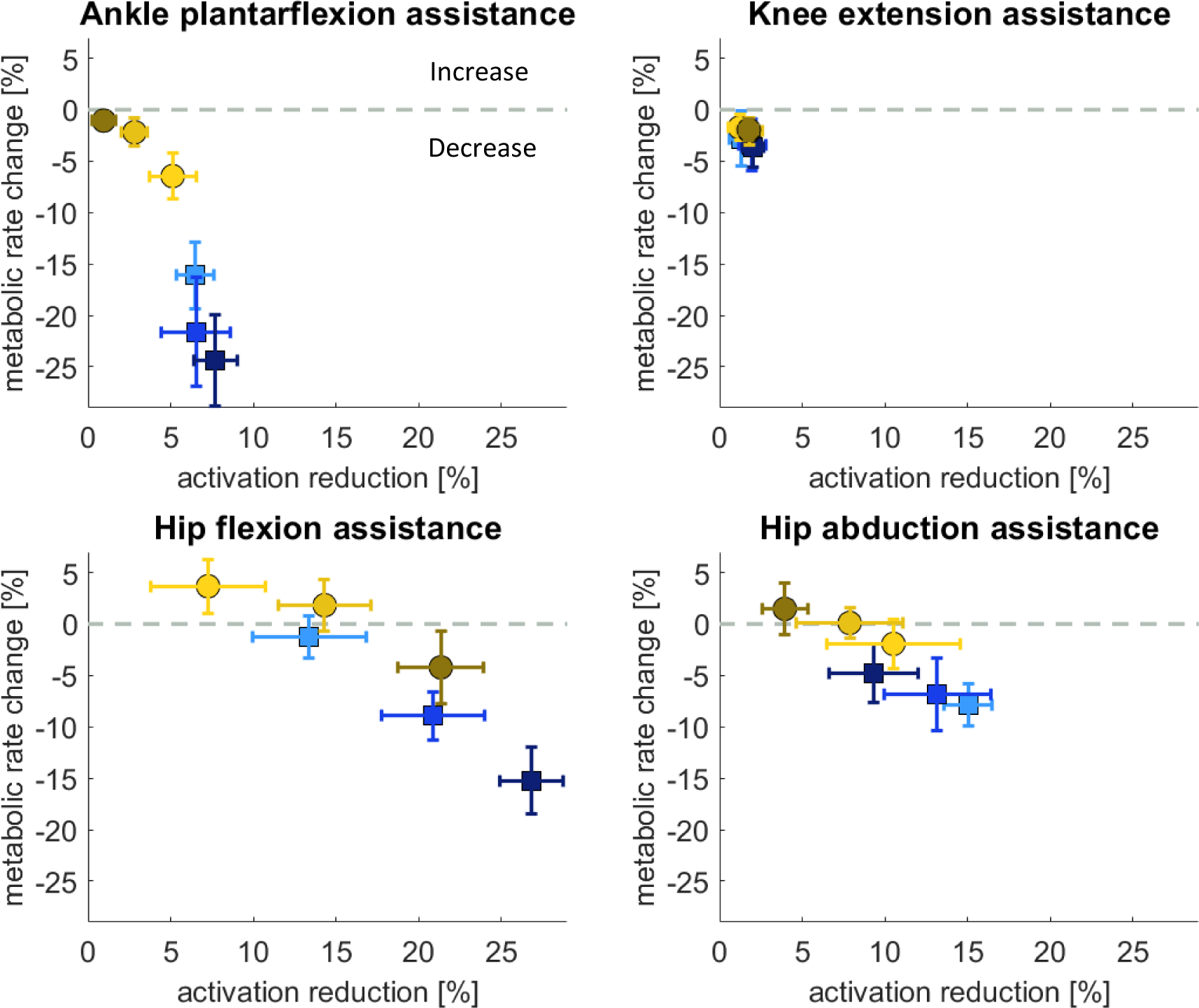
Change in metabolic rates vs. reduction of muscle activations, shown as % of unassisted conditions, at slow, normal, and fast walking speeds with motor-based and spring-based assistance. The values shown are average ± 1 standard deviation among all subjects and gait cycles.

With spring-based actuation, relative muscle activations were nearly with identical trends as with motor-based actuation, though all proportionally lower, with one major contrast, that muscle activation changes with plantarflexion assistance were inversely proportional to walking speed, and were practically zero at fast walking speed.

While relative muscle activation changes were largely proportional to relative metabolic rate changes, they did not always translate to reduced metabolic cost; spring-based assistance actually resulted in 2-4% higher metabolic rates, most notably with hip flexion assistance at slow and normal speeds and with hip abduction assistance at fast speed. The largest reduction (average ca. 7%) of relative metabolic rate with spring-based actuation resulted from ankle plantarflexion assistance at slow speed, followed ca. 5% reduction with hip flexion reduction at fast speed.

Motor-based assistance always caused a decrease in metabolic rates, wherein the highest relative reduction (average ca. 24%) was observed with ankle plantarflexion assistance at fast speed, followed by ankle plantarflexion assistance at lower speeds (22% at normal and 16% at slow speeds) then by hip flexion assistance (15%) at high walking speed. Hip flexion assistance at low speed had practically no effect on metabolic rate change, nor did knee extension assistance at any speed.

Analyses of the influence of ideal assistive moments at each joint are described in more detail in the next section.

### 3.2. Ankle plantarflexion assistance

The computed ideal motor-based plantarflexion assistance contributed with more than half of the net ankle plantarflexion moment, and only increased slightly in magnitude with increasing speed; the net plantarflexion muscle moment was reduced by approximately 60% at all speeds (Fig. 3), while the net dorsiflexor muscle moment increased by up to 4%. With motor-based assistance, the total metabolic rate peak at all speeds was reduced near terminal stance and pre-swing phases. Overall, these differences resulted in a 16% reduction in overall metabolic rate in slow walking and a 24% reduction in fast walking. Soleus activation was nearly entirely reduced with motor-based plantarflexion assistance, and, to a lower extent, gastrocnemius activation (Supplementary Fig. 2). The gastrocnemius still generated a moment during midstance, contributing to the ankle plantarflexion and knee flexion moments. Tibialis anterior activation remains nearly the same compared to unassisted conditions during mid-stance.

**Fig. 3.**
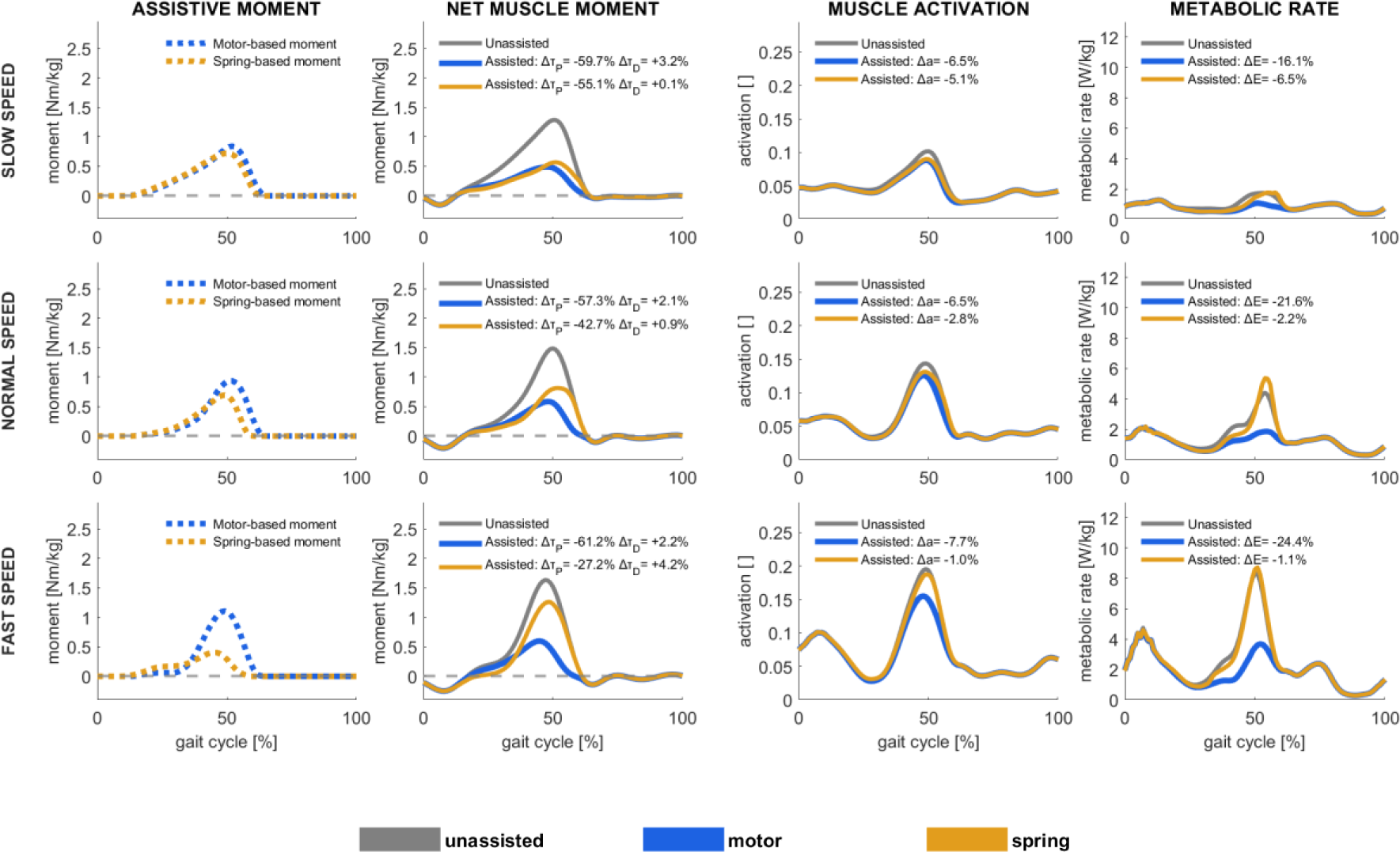
Assistive device moments [first column], net muscle moments [second column], muscle activations [third column], and metabolic rates [fourth column] in unassisted conditions and with motor-based and spring-based assistance during slow (upper row), normal (middle row), and fast (lower row) walking speed, shown as average values among all subjects and gait cycles. Positive moment refers to ankle plantarflexion, and negative to ankle dorsiflexion. Change in ankle plantarflexion (*Δτ*_*P*_) and ankle dorsiflexion (Δτ_D_) moments, muscle activations (Δa), and metabolic rates (ΔE), shown as % of unassisted conditions, are presented.

Ideal spring-based plantarflexion assistance contributed with more overall moments in slow walking than in normal or fast walking; the plantarflexor muscle moment was reduced by more than half (55%) in slow walking, by 43% in normal and 27% in fast walking. With spring-based assistance, the total metabolic rate peak was reduced by 7% in slow walking, 2% in normal, and 1% in fast walking. The peak ankle dorsiflexion angle, which sets the assistive moment peak, occurs earlier in the gait cycle as walking speed increases; the spring can thus not maximally assist the muscle plantarflexor moment peak at pre-swing to the same extent as motor-based actuation can. During terminal stance, soleus and gastrocnemius activations were reduced with spring-based assistance, but tibialis anterior activations were increased. Muscle fiber velocities increased in the soleus and gastrocnemius during push-off, and, as a result, muscle positive power increased (supplementary Fig. 2 and 3), resulting in increased total metabolic rate peak at all speeds even though the average metabolic rate over the gait cycle decreased (supplementary Fig. 4).

### 3.3. Knee extensor assistance

Ideal motor-based knee extensor assistance was only effectual in loading response and early midstance, where it contributed with nearly all knee extensor moments at all walking speeds (Fig. 4). The assistive moment resulted in a net muscle moment decrease of 47-50% at all speeds. The assistive moment resulted in a slightly increased knee flexion moment just after initial contact, more so at high walking speed. With assistance, during loading response, vasti activations decreased, but muscle power increased (supplementary Fig. 2 and 3); knee extension assistance resulted in decreased vasti tendon force, which decreased tendon strain and thus increased fiber velocities. As a result, both muscle negative power during loading response and muscle positive power in early midstance increased. Consequently, metabolic rates from vasti dynamics decreased in loading response and increased slightly in early midstance (supplementary Fig. 4). Overall, the motor-based assistance resulted in a 2-3% metabolic rate reduction at all walking speeds.

**Fig. 4.**
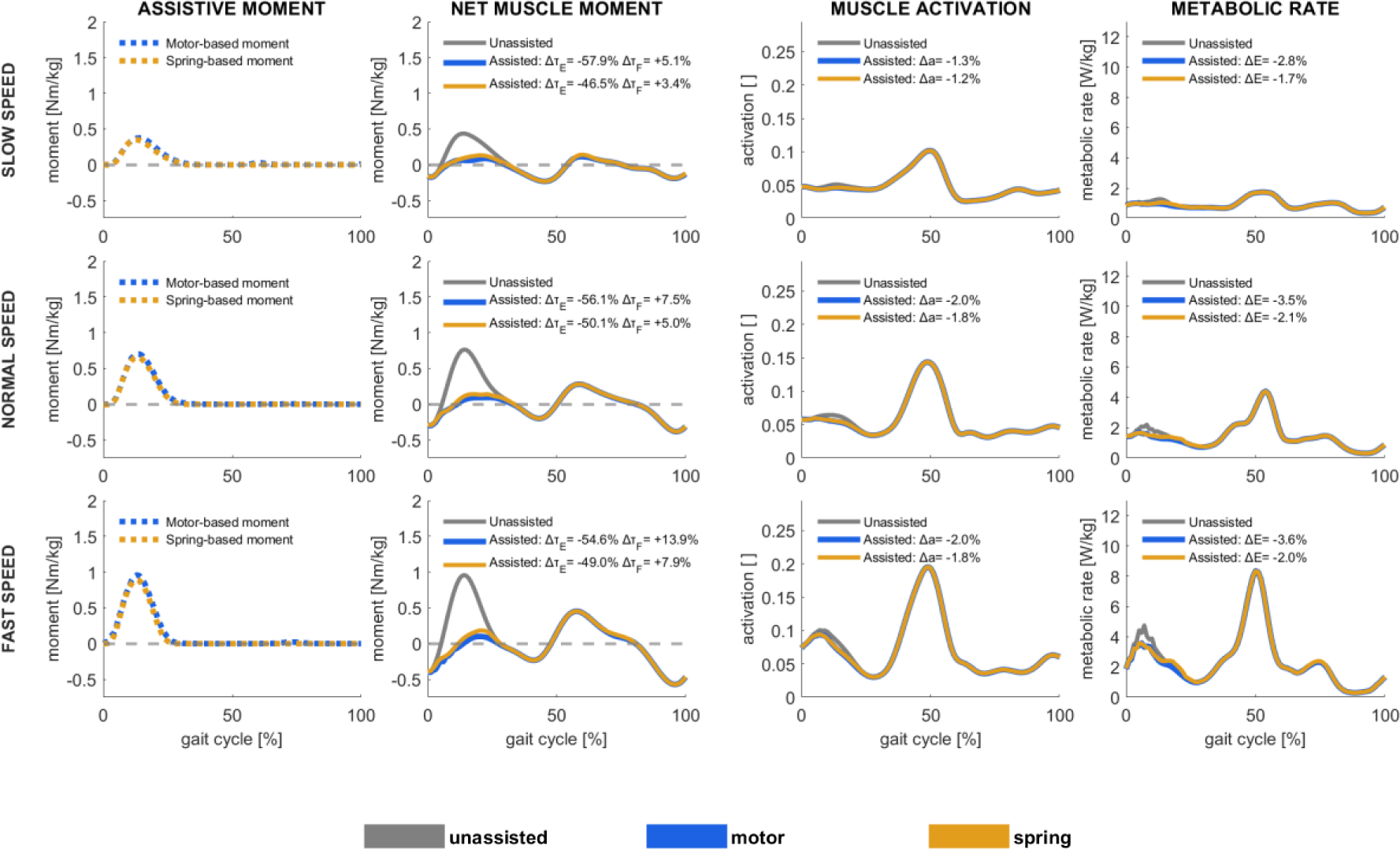
Assistive device moments [first column], net muscle moments [second column], muscle activations [third column], and metabolic rates [fourth column] in unassisted conditions and with motor-based and spring-based assistance during slow (upper row), normal (middle row), and fast (lower row) walking speed, shown as average values among all subjects and gait cycles. Positive moment refers to knee extension, and negative to knee flexion. Change in knee extension (*Δτ*_*E*_) and knee flexion (Δτ_F_) moments, muscle activations (Δa), and metabolic rates (ΔE), shown as % of unassisted conditions, are presented.

Ideal spring-based knee extensor assistance was likewise only effectual in loading response and early midstance, to practically the same degree as motor-based assistance. It resulted in similar reductions in muscle activations, net muscle moments, and metabolic energy rates, yet to a somewhat lower magnitude; with assistance, the total metabolic rate was reduced by approximately 2% at all speeds.

### 3.4. Hip flexor assistance

Ideal motor-based hip flexor assistance was effectual largely in terminal stance and preswing, increasing with walking speed, and mid- to late swing (Fig. 5) and to a very small amount immediately after initial contact. The assistive moment resulted in substantially decreased hip flexion muscle moment, ranging from 66% reduction at slow and 80% at fast walking speeds, mostly observed in terminal stance and preswing, but also *increased* hip extensor muscle moment in mid- to late swing. The increase in hip extensor muscle moment was relatively similar at all speeds but led to a particularly remarkable 168% increase in net hip extensor muscle moment in slow walking, during which the extensor moment was negligible without assistance. The increase in hip extension muscle moment reflects a trade-off between decreased activations in the hip flexion muscle group (see psoas in supplementary Fig. 2) at the expense of slightly increased activations in other muscle groups (see biceps femoris long head and vastus lateralis in supplementary Fig. 2). As a result, with assistance, metabolic rates were reduced during terminal stance and pre-swing but increased during early to mid-swing. Overall, with motor-based hip flexor assistance, the total metabolic rate decreased by 15% in fast walking, 9% in normal and 1% in slow walking. Without assistance, the vasti were most active during loading response and mid-stance, but with motor-based assistance, the vasti were also active during mid-swing, likely as antagonists for the increased biceps femoris long head activation. This activation pattern resulted in increased vasti force and power during the swing phase (supplementary Fig. 3), which caused vasti negative power during initial swing and positive power during mid-swing. As muscle positive power is associated with higher metabolic rates, motor-based assistance resulted in slightly increased metabolic rates during mid-swing.

**Fig. 5.**
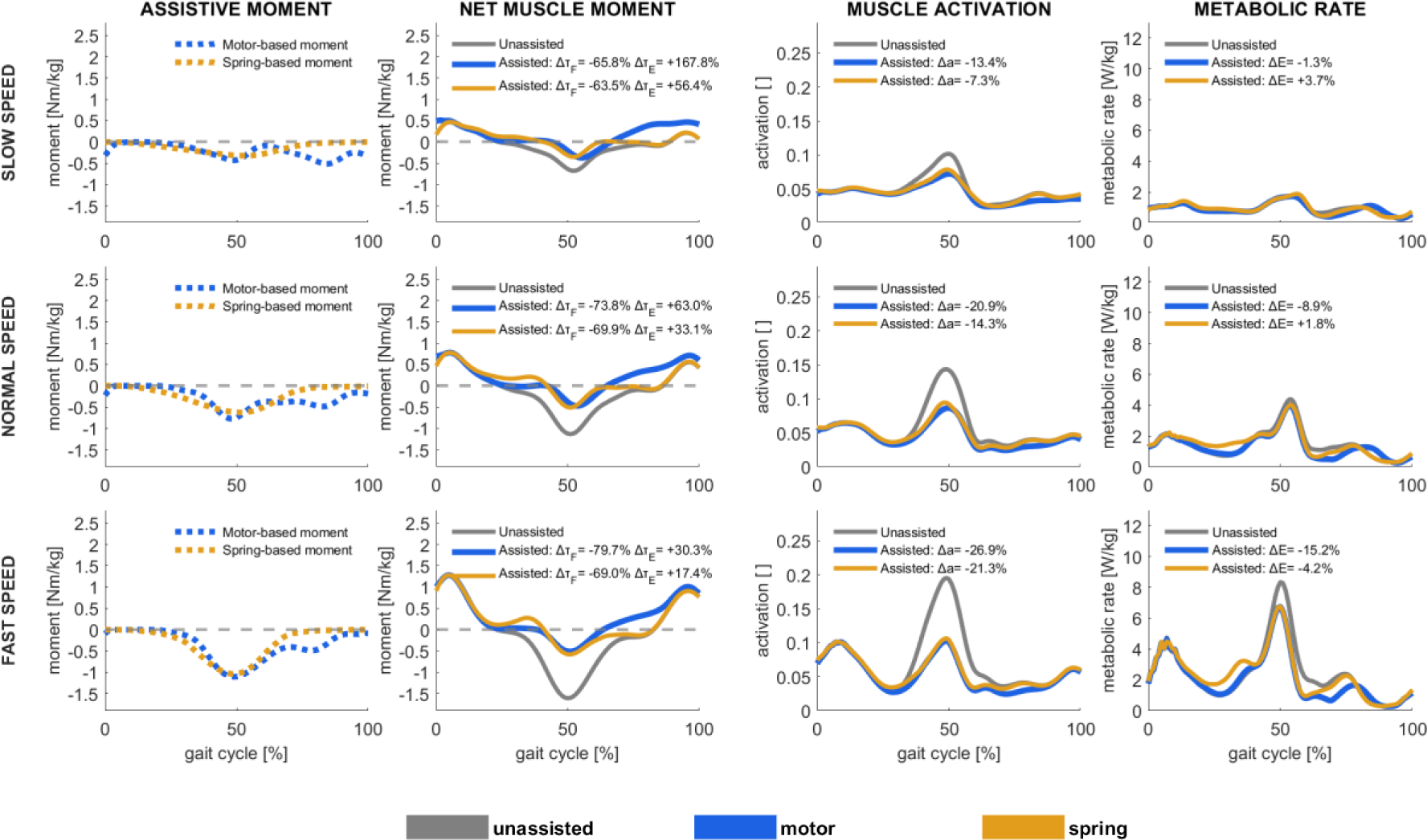
Assistive device moments [first column], net muscle moments [second column], muscle activations [third column], and metabolic rates [fourth column] in unassisted conditions and with motor-based and spring-based assistance during slow (upper row), normal (middle row), and fast (lower row) walking speed, shown as average values among all subjects and gait cycles. Positive moment refers to hip extension, and negative to hip flexion. Change in hip extension (*Δτ*_*E*_) and hip flexion (Δτ_F_) moments, muscle activations (Δa), and metabolic rates (ΔE), shown as % of unassisted conditions, are presented.

Ideal spring-based hip flexor assistance was only effectual during terminal stance and preswing, as it is set by spring engagement as the hip extends during mid-stance and disengagement as the hip flexes in early swing (Fig. 5). With assistance, the hip flexor muscle moment was greatly reduced during this phase; the net hip flexor muscle moment was reduced by 64 in slow and 69-70% in faster walking. However, its engagement during midstance, which accommodated energy storage during hip extension, resulted in *increased* hip extensor muscle during midstance. With assistance, the gluteus maximum and semimembranosus activations increased in midstance, and vasti activation increased in initial swing (Supplementary Fig. 2), resulting in higher muscle positive power and, thereby, metabolic rates during the mid- to terminal stance. In contrast, the increased vasti activation corresponded to higher muscle negative power, which did not increase metabolic rates (supplementary Fig. 3 and 4). Overall, with spring-based hip flexor assistance, the total metabolic rate decreased only during fast walking (4%) but increased by 2-4% in normal and slow walking.

### 3.5. Hip abduction assistance

Ideal motor-based hip abduction assistance was effectual throughout nearly the entire stance phase, accounting for the majority of net hip abduction moment, reducing the hip abductor muscle moment by more than 70% at all walking speeds and more at slower speeds (Fig. 6). The assistive moment peaked at approximately 20 and 50% of the gait cycle. Whereas the first assistive peak reduced the net muscle hip abduction moments and hip abductor muscle activations, the second peak increased the net hip adduction moment and adductor muscle activations (Supplementary Fig. 2), with correspondingly higher hip adductor muscle positive power and metabolic rates (supplementary Fig. 3 and 4). Overall, with motor-based hip abduction assistance, the total metabolic rate decreased by 7-8% in normal and fast walking and by 5% in slow walking.

**Fig. 6.**
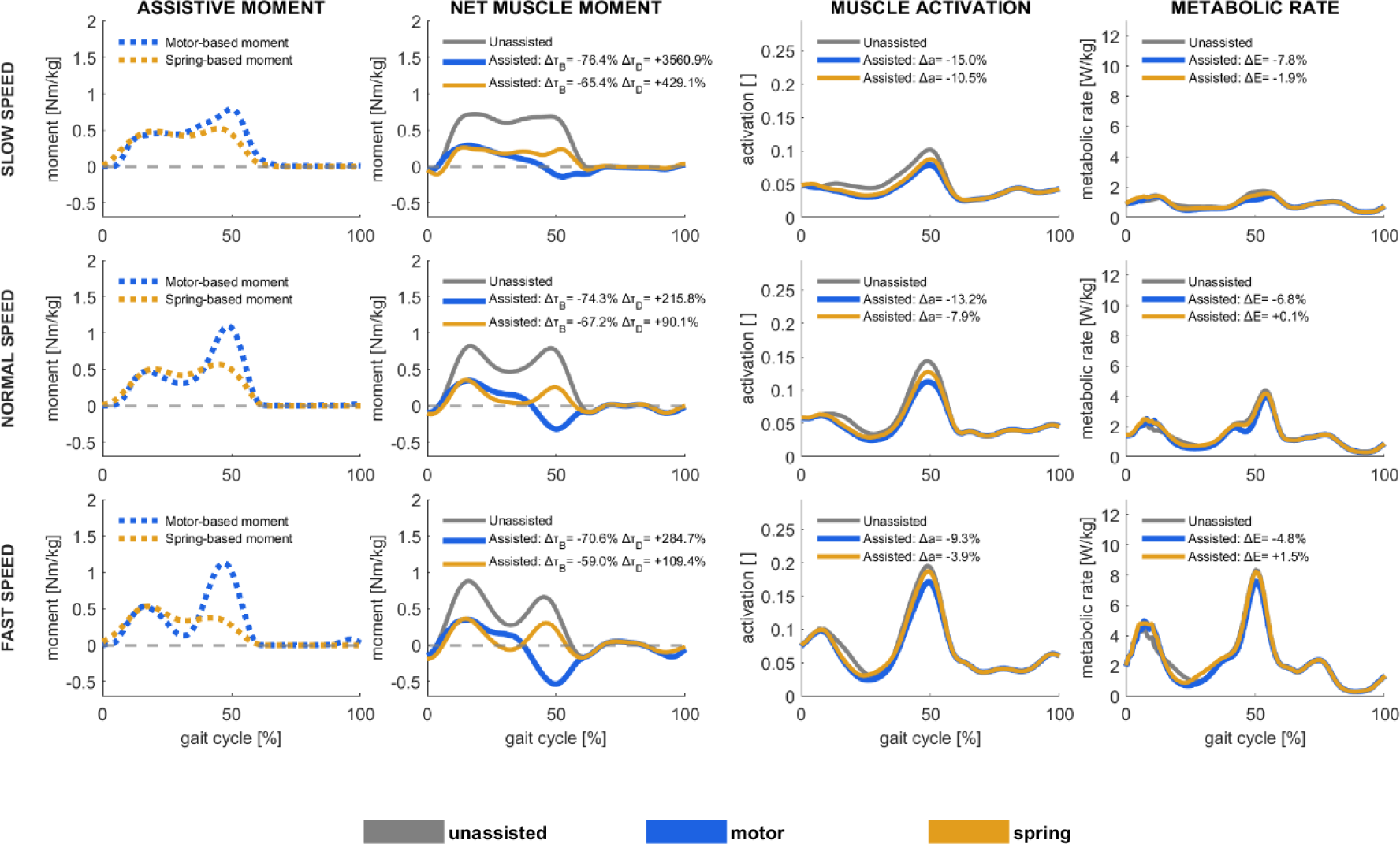
Assistive device moments [first column], net muscle moments [second column], muscle activations [third column], and metabolic rates [fourth column] in unassisted conditions and with motor-based and spring-based assistance during slow (upper row), normal (middle row), and fast (lower row) walking speed, shown as average values among all subjects and gait cycles. Positive moment refers to hip abduction, and negative to hip adduction. Change in hip abduction (*Δτ*_*B*_) and hip adduction (Δτ_D_) moments, muscle activations (Δa), and metabolic rates (ΔE), shown as % of unassisted conditions, are presented.

Ideal spring-based hip abduction assistance was likewise effectual during nearly the entire stance phase. With spring-based assistance, the hip abductor muscle moment decreased by approximately 60% at all walking speeds. However, the overall metabolic rate was nearly unchanged; with assistance, the metabolic rate decreased by 2% in slow walking, was unchanged in normal walking, and increased by 2% in fast walking. The spring-based assistance had a less pronounced peak in terminal stance than motor-based assistance, as it was set by the hip adduction angle, and a hip abductor muscle moment was still required in this phase, though lower than without assistance. Similar to motor-based assistance, spring-based assistance involved a trade-off between decreased hip abductor muscle activation and increased hip adductor muscle activation (supplementary Fig. 2). This trade-off was, however, even less effective in reducing activations and metabolic rates than the motor-based assistance. While metabolic rates decreased in gluteus medialis and minimus, and tensor fasciae latae with spring-based assistance, they did not decrease as much as with motor-based assistance. Also, metabolic rates in the gluteus maximum during mid-stance were higher with spring-based than with motor-based assistance.

## 4. Discussion

In this simulation study, ideal assistive moments were identified, defined as those that reduced the squared sum of muscle activations. The assistive moment profiles in a motor-based actuator could have a variable profile, but those with the spring-based actuators were constrained by joint kinematics. The ideal assistive moments in both actuator modes substantially decreased net muscle moments, i.e., the net joint moment minus the assistive moment. Whereas motor-based assistance always reduced total metabolic rates to some extent, varying among joints and speeds, spring-based assistance did not always reduce metabolic rates. The most notable reductions in metabolic rates resulted from motor-based plantarflexion assistance, followed by motor-based hip flexion assistance, both more effective at higher speeds. Motor-based hip abduction assistance also reduced metabolic rate, interestingly inversely with walking speed. Spring-based hip flexion assistance at slow and normal speeds and hip abduction assistance at normal and fast speeds reduced muscle activations to some extent, but these reductions did not translate to reduced metabolic rates; rates were unchanged or even increased slightly. Knee extension assistance, regardless of actuation mode or walking speed, had little to no effect on metabolic rates, even though it was able to contribute to a majority of the net extensor moment in loading response.

Our findings indicate that an assistive strategy based on minimal muscle activations does not translate to a decreased metabolic rate. Assistive devices are generally designed to support motion, which might involve reducing net muscle moment, activations, and metabolic rates (2). The optimal assistive moments in our simulations decreased the overall sum of muscle activation, which in turn reduced muscle forces and, thus, net muscle moment in the assisted muscle groups, though occasionally increasing demand on antagonist muscles. Reduction of metabolic rates was, however, more difficult to achieve. Metabolic energy models estimate muscle energy rates based on heat dissipation and muscle power. Heat dissipation is the sum of various subcomponents that depend on fiber velocities, such as the shortening and lengthening heat rates and the activation and maintenance heat rates (30). All components are related to muscle activations. As such, metabolic rate is diminished if heat dissipation and muscle power are zero, which only happens when muscle activations are zero. However, assistive moments that submaximally reduce muscle activations, i.e. lower but non-zero activations, can result in higher metabolic rates if the muscle positive power increase outweighs the heat dissipation decrease. We observed two instances in which this was the case, both with spring-based assistance: With plantarflexion assistance during preswing, and with knee extension assistance during mid-stance, wherein lower activation was associated with higher fiber velocity in the assisted muscles, which increased muscle positive power, and thereby metabolic rates. In several cases, assistive moment resulted in increased demand, and thus muscle positive power and metabolic rates, in antagonist muscles, for instance, with hip flexion assistance during mid-swing and with hip abduction assistance during loading response. It is not a straightforward assumption that assistive moments that reduce overall muscle activations will also reduce metabolic rates. We did, however, identify several cases in which the assistive moment reduced agonist muscle activations to zero, without substantially increasing antagonist muscle activations, and resulted in overall reduced metabolic rates, specifically with motor-based ankle plantarflexion and knee extension assistance.

Our identified ideal motor-based plantarflexion assistive moment profiles are similar to previously reported moment profiles that were found to reduce metabolic rates, but our ideal spring-based plantarflexion assistive moments disagreed somewhat. At normal walking speed, the ideal motor-based plantarflexion assistive moment profile was similar to those identified from human-the-loop optimization studies that aimed for minimal metabolic rates (3,31). The peak in the profile we identified was near 50% of the gait cycle, agreeing with other identified optimal moment trajectories (3,31). However, our simulation predicts a metabolic reduction (22%) at normal walking speed, which is substantially larger than experimentally reported metabolic rate reduction reported by Zhang et al. at a slightly lower speed (14% metabolic reduction at 1.25 m/s) (31). We also found that with motor-based plantarflexion assistance, the metabolic reduction should be more pronounced as walking speed increases, in agreement with prior studies (3,31). However, experimental comparisons with spring-based assistance are more challenging, as very few studies have studied muscle activation changes across walking speeds. In a study from Nuckols and Sawicki using an exoskeleton emulator to mimic spring-like actuation, the optimal spring stiffness to reduce metabolic rates was similar at speeds of 1.25 and 1.75 m/s (32). This finding does not align with our results, presumably because our optimal assistive moment does not directly minimize metabolic rates.

Our findings of decreased muscle activation but increased muscle positive power during loading response with knee extension assistance might explain why previous experimental studies with this aim have failed to reduce metabolic rates during walking. Metabolic rate reduction with knee extension assistance has only been achieved with motor-based actuation compared to wearing a powered-off exoskeleton (33,34) or in challenging environments such as carrying loads while walking on an inclined surface (35). Our simulation suggests that, with knee extensor assistance, the knee extensor muscle forces decrease during loading response, resulting in decreased tendon strain and, thus, higher muscle fiber velocities and muscle positive power, which in turn actually increased the muscle metabolic rates. Jackson et al. reported a similar finding that even if muscle moment is reduced with an ankle exoskeleton, the metabolic cost can increase if the muscle moment corresponds to increasing muscle positive work (7). In our study, we found little to no potential benefit from knee extensor assistance, regardless of actuation mode or walking speed.

Spring-based and especially motor-based hip flexion assistance shows promise in reducing metabolic rates, particularly at fast walking speeds. Only a few studies have evaluated the effects of hip flexion assistance with powered devices (36–38). Studies of devices that reduced metabolic cost parametrized the assistive profile such that it began at maximum hip extension (38) or provided a power burst during a predefined time window (corresponding to 25% of the gait cycle) (37); both studies found that the optimal peak assistive moment should be later than the net hip flexion moment. These assistive trajectories disagree somewhat with the optimal assistive moments that we identified. We also found that the increased antagonist hip extensor muscle moment during the swing phase counterintuitively reduced metabolic rate, in agreement with findings reported in another simulation study (9). It is possible that this counterintuitive benefit from simulation studies might not translate experimentally; it has, to the best of our knowledge, not been tested experimentally. Furthermore, previous experimental studies reported a metabolic decrease of 8.8% with hip flexion assistance compared to unassisted conditions (38) and 6.1% compared with a powered-off exoskeleton (39), which agrees with our predictions (9% near preferred walking speed). With spring-based hip flexion assistance, only two experimental studies reported metabolic rate reduction (19,40). Zhou et al. reported 7.2% metabolic rate reduction at 1.5 m/s and suggested that optimal assistive is likely speed-dependent (19). We found no metabolic reduction with spring-based assistance at preferred walking speed but a small decrease (4%) in fast walking. To the best of our knowledge, no previous study has evaluated the influence of hip flexion assistance, neither spring- or motor-based, at different walking speeds. Our simulation supports a hypothesis that hip flexion assistance from either actuation mode can potentially decrease metabolic rate as walking speed increases.

Our findings indicate little to no potential for hip abduction assistance to substantially reduce metabolic rates. Only one recent pilot experimental study has evaluated metabolic rates with hip abduction assistance (41). Kim et al. found that human-in-the-loop optimization with a motor-based actuation did not reduce metabolic rates. They attributed this finding to the role of the hip abductors during walking, which stabilizes the hip and maintains balance, and suggested that minimizing their muscle activity may not be an advantageous strategy for metabolic rate reduction. Our findings likewise suggest that, as speed varies, preserving balance remains the dominant objective of hip abductor activation, as metabolic rate changes are inversely proportional to speed.

The two major limitations of our study are 1) the assumption that motion patterns in unassisted and assisted conditions are unchanged, which is a dilemma in all musculoskeletal simulations with constrained kinematics, and 2) the optimal assistive moments defined with the objective function in the muscle redundancy solver that seeks the task-specific exoskeleton moments that minimize the sum of squared muscle activations. The assumption of unchanged kinematics might be reasonable in spring-based devices at the ankle (18) and hip (19), as they can be made lightweight and reasonably comfortable. With powered ankle and hip assistive devices, despite evidence suggesting that joint angles and net joint moments might be preserved (36,42), human-device adaptation is complex and more likely to alter the user’s motor control strategy and, thereby, joint kinematics and moments (43). Regarding the second limitation, we formulated the optimization problem to solve muscle redundancy and to identify optimal assistive moments using the same objective function, specifically minimal muscle activation. We assumed the paradigm that human walking is achieved by minimal muscle activations, and that metabolic efficiency is driven by this neuromuscular strategy. As such the assistive moments in our simulation represent the optimal for minimal muscle activations, but we demonstrated that minimal activations and metabolic rates are not necessarily aligned. Our findings warrant further simulation studies that identify assistive moments with optimization goals other than minimal activations, ideally with goals of minimal metabolic cost. In this regard, it might be beneficial to explicitly incorporate the goal of the assistive moment separately from the objective function to solve muscle redundancy. This can theoretically be achieved by adopting a bilevel optimization scheme as proposed by Nguyen et al. (44), in which a low level optimization problem might deal with solving muscle redundancy, while a upper level problem searches for optimal assistance with a task criterion e.g., maximal walking stability or minimal metabolic cost. Furthermore, forward dynamics simulation studies with assistive devices have great potential to predict musculoskeletal skeletal dynamics and to explicitly formulate the goals with assistive devices.

## 5. Data availability

Experimental data to replicate this study, such as subject anthropometrics, marker trajectories, and ground reaction forces are available in the following repository: https://figshare.com/s/1caa0e14c79426cb12cc

## 6. Code availability

The scripts for simulating exoskeleton assistance with tuned muscle-tendon parameters and computing metabolic rates based on the metabolic energy models are available in the following repository: https://github.com/israelluis/Exoskeletons_ExperimentGuidedCalibration

## 7. Author contributions

**Israel Luis**: Conceptualization, Software, Formal analysis, Writing – Original Draft Preparation. **Maarten Afschrift**: Software, Formal analysis and Writing - Review & Editing. **Elena M. Gutierrez-Farewik**: Formal analysis, Writing - Review & Editing and Supervision.

## 8. Competing interests

The authors declare no competing interests.

## 9. Materials & Correspondence

Correspondence and requests for materials should be addressed to Israel Luis.

## 10. Acknowledgment

Authors acknowledge the funding sources provided by the Swedish Research Council (nr 2018-00750) and Promobilia Foundation (nr 18200).

## Notes

### Competing Interest Statement

The authors have declared no competing interest.

## Bibliography

1. Sawicki GS, Beck ON, Kang I, Young AJ. The exoskeleton expansion: Improving walking and running economy. J Neuroeng Rehabil. 2020;17(1):1–9.

2. Young AJ, Ferris DP. State-of-the-art and Future Directions for Robotic Lower Limb Exoskeletons. IEEE Transactions on Neural Systems and Rehabilitation Engineering. 2016;PP(99):1–1.

3. Slade P, Kochenderfer MJ, Delp SL, Collins SH. Personalizing exoskeleton assistance while walking in the real world. Nature. 2022 Oct 13;610(7931):277–82.

4. Kapelner T, Sartori M, Negro F, Farina D. Neuro-Musculoskeletal Mapping for Man-Machine Interfacing. Scientific Reports. 2020;10(1):1–10.

5. Liu YX, Gutierrez-Farewik EM. Joint Kinematics, Kinetics and Muscle Synergy Patterns During Transitions Between Locomotion Modes. IEEE Transactions on Biomedical Engineering. 2023;70(3):1062–71.

6. Grabke EP, Masani K, Andrysek J. Lower Limb Assistive Device Design Optimization Using Musculoskeletal Modeling: A Review. Journal of Medical Devices, Transactions of the ASME. 2019;13(4):1–13.

7. Jackson RW, Dembia CL, Delp SL, Collins SH. Muscle-tendon mechanics explain unexpected effects of exoskeleton assistance on metabolic rate during walking. Journal of Experimental Biology. 2017;220(11):2082–95.

8. Sawicki GS, Khan NS. A Simple Model to Estimate Plantarflexor Muscle-Tendon Mechanics and Energetics During Walking With Elastic Ankle Exoskeletons. IEEE Transactions on Biomedical Engineering. 2016;63(5):914–23.

9. Dembia CL, Silder A, Uchida TK, Hicks JL, Delp SL. Simulating ideal assistive devices to reduce the metabolic cost of walking with heavy loads. PLoS One. 2017;12(7):1–25.

10. Cseke B, Uchida TK, Doumit M. Simulating Ideal Assistive Strategies to Reduce the Metabolic Cost of Walking in the Elderly. IEEE Trans Biomed Eng. 2022;69(9):2797–805.

11. Uchida TK, Seth A, Pouya S, Dembia CL, Hicks JL, Delp SL. Simulating ideal assistive devices to reduce the metabolic cost of running. PLoS One. 2016;11(9):1–19.

12. Franks PW, Bianco NA, Bryan GM, Hicks JL, Delp SL, Collins SH. Testing Simulated Assistance Strategies on a Hip-Knee-Ankle Exoskeleton: A Case Study. Proceedings of the IEEE RAS and EMBS International Conference on Biomedical Robotics and Biomechatronics. 2020;2020-Novem:700–7.

13. Lee G, Kim J, Panizzolo FA, Zhou YM, Baker LM, Galiana I, et al. Reducing the metabolic cost of running with a tethered soft exosuit. Sci Robot. 2017 May 31;2(6):1–3.

14. Perry Jacquelin. Gait analysis : normal and pathological function. Second Edition. 1992. 1–576 p.

15. Van Dijk W, Van Der Kooij H, Hekman E. A passive exoskeleton with artificial tendons: Design and experimental evaluation. IEEE International Conference on Rehabilitation Robotics. 2011;

16. Sawicki GS, Khan NS. A Simple Model to Estimate Plantarflexor Muscle-Tendon Mechanics and Energetics During Walking With Elastic Ankle Exoskeletons. IEEE Trans Biomed Eng. 2016 May 1;63(5):914–23.

17. Chen W, Wu S, Zhou T, Xiong C. On the biological mechanics and energetics of the hip joint muscle-tendon system assisted by passive hip exoskeleton. Bioinspir Biomim. 2019 Jan 1;14(1).

18. Collins SH, Wiggin MB, Sawicki GS. Reducing the energy cost of human walking using an unpowered exoskeleton. Nature. 2015;522(7555):212–5.

19. Zhou T, Xiong C, Zhang J, Hu D, Chen W, Huang X. Reducing the metabolic energy of walking and running using an unpowered hip exoskeleton. Journal of NeuroEngineering and Rehabilitation. 2021;18(1):1–15.

20. Luis I, Afschrift M, De Groote F, Gutierrez-Farewik EM. Insights into muscle metabolic energetics: Modelling muscle-tendon mechanics and metabolic rates during walking across speeds. 2023;

21. Seth A, Hicks JL, Uchida TK, Habib A, Dembia CL, Dunne JJ, et al. OpenSim: Simulating musculoskeletal dynamics and neuromuscular control to study human and animal movement. Schneidman D, editor. PLoS Comput Biol. 2018;14(7):e1006223.

22. Luis I, Afschrift M, Gutierrez-Farewik EM. Experiment-Guided Calibration of Muscle Fiber Lengths and Passive Forces. 2023;

23. De Groote F, Kinney AL, Rao A V., Fregly BJ. Evaluation of Direct Collocation Optimal Control Problem Formulations for Solving the Muscle Redundancy Problem. Ann Biomed Eng. 2016;44(10):2922–36.

24. Rajagopal A, Dembia CL, DeMers MS, Delp DD, Hicks JL, Delp SL. Full-Body Musculoskeletal Model for Muscle-Driven Simulation of Human Gait. IEEE Trans Biomed Eng. 2016;63(10):2068–79.

25. Uhlrich SD, Jackson RW, Seth A, Kolesar JA, Delp SL. Muscle coordination retraining inspired by musculoskeletal simulations reduces knee contact force. Scientific Reports. 2022;12(1):1–13.

26. Arnold EM, Hamner SR, Seth A, Millard M, Delp SL. How muscle fiber lengths and velocities affect muscle force generation as humans walk and run at different speeds. Journal of Experimental Biology. 2013;216(11):2150–60.

27. Farris DJ, Raiteri BJ. Elastic ankle muscle-tendon interactions are adjusted to produce acceleration during walking in humans. Journal of Experimental Biology. 2017;220(22):4252–60.

28. Bohm S, Marzilger R, Mersmann F, Santuz A, Arampatzis A. Operating length and velocity of human vastus lateralis muscle during walking and running. Sci Rep. 2018;8(1):1–10.

29. Silder A, Whittington B, Heiderscheit B, Thelen DG. Identification of passive elastic joint moment-angle relationships in the lower extremity. J Biomech. 2007;40(12):2628–35.

30. Bhargava LJ, Pandy MG, Anderson FC. A phenomenological model for estimating metabolic energy consumption in muscle contraction. J Biomech. 2004;37(1):81–8.

31. Zhang J, Fiers P, Witte KA, Jackson RW, Poggensee KL, Atkeson CG, et al. Human-in-the-loop optimization of exoskeleton assistance during walking. Science. 2017 Jun 23;356(6344):1280–4.

32. Nuckols RW, Nuckols RW, Nuckols RW, Sawicki GS, Sawicki GS. Impact of elastic ankle exoskeleton stiffness on neuromechanics and energetics of human walking across multiple speeds. J Neuroeng Rehabil. 2020;17(1):1–19.

33. Zhou Z, Liao Y, Wang C, Wang Q. Preliminary evaluation of gait assistance during treadmill walking with a light-weight bionic knee exoskeleton. In: 2016 IEEE International Conference on Robotics and Biomimetics, ROBIO 2016. Institute of Electrical and Electronics Engineers Inc.; 2016. p. 1173–8.

34. Franks PW, Bryan GM, Martin RM, Reyes R, Lakmazaheri AC, Collins SH. Comparing optimized exoskeleton assistance of the hip, knee, and ankle in single and multi-joint configurations. Wearable Technologies. 2021 Nov 24;2:e16.

35. MacLean MK, Ferris DP. Energetics of walking with a robotic knee exoskeleton. Journal of Applied Biomechanics. 2019;35(5):320–6.

36. Lewis CL, Ferris DP. Invariant hip moment pattern while walking with a robotic hip exoskeleton. J Biomech. 2011 Mar 15;44(5):789–93.

37. Young AJ, Foss J, Gannon H, Ferris DP. Influence of power delivery timing on the energetics and biomechanics of humans wearing a hip exoskeleton. Front Bioeng Biotechnol. 2017 Mar 8;5(MAR).

38. Kim J, Quinlivan BT, Deprey LA, Arumukhom Revi D, Eckert-Erdheim A, Murphy P, et al. Reducing the energy cost of walking with low assistance levels through optimized hip flexion assistance from a soft exosuit. Sci Rep. 2022 Dec 1;12(1).

39. Young AJ, Foss J, Gannon H, Ferris DP. Influence of power delivery timing on the energetics and biomechanics of humans wearing a hip exoskeleton. Frontiers in Bioengineering and Biotechnology. 2017;5(MAR):1–11.

40. Panizzolo FA, Bolgiani C, Di Liddo L, Annese E, Marcolin G. Reducing the energy cost of walking in older adults using a passive hip flexion device. Journal of NeuroEngineering and Rehabilitation. 2019;16(1):1–9.

41. Kim J, Raitor M, Liu CK, Collins SH. Frontal hip exoskeleton assistance does not appear promising for reducing the metabolic cost of walking: A preliminary experimental study. 2023;

42. Gordon KE, Ferris DP. Learning to walk with a robotic ankle exoskeleton. J Biomech. 2007;40(12):2636–44.

43. Poggensee KL, Collins SH. How adaptation, training, and customization contribute to benefits from exoskeleton assistance. Sci Robot. 2021;6(58):1–44.

44. Nguyen VQ, Johnson RT, Sup FC, Umberger BR. Bilevel Optimization for Cost Function Determination in Dynamic Simulation of Human Gait. IEEE Transactions on Neural Systems and Rehabilitation Engineering. 2019;27(7):1426–35.

